# EMG-BIDS: an extension to the Brain Imaging Data Structure for electromyography

**DOI:** 10.64898/2026.07.21.739911

**Authors:** Seyed Yahya Shirazi, Daniel McCloy, Robert Oostenveld, Tjeerd Boonstra, Eric Larson, Julius Welzel, Remi Gau, Christopher J. Markiewicz, Simone Posella, Jörn M. Horschig, Thomas Klotz, Alexandre Gramfort, Arnaud Delorme

## Abstract

Electromyography (EMG) is fundamental to clinical assessment, rehabilitation, neuromuscular research, and human-machine interfaces. Despite decades of use, no widely adopted standard exists for organizing and sharing EMG data, limiting reusability and large-scale data aggregation. We present EMG-BIDS, an extension to the Brain Imaging Data Structure (BIDS) that standardizes the organization of EMG recordings. EMG-BIDS addresses challenges unique to EMG, including diverse electrode types (surface or intramuscular, single channel to high-density arrays), heterogeneous electrode placements across anatomical locations, montages (e.g., monopolar or bipolar sensor designs), and the critical need for transparent documentation of sensor positioning. The specification introduces hierarchical coordinate systems that link local electrode grids to anatomical landmarks, enabling precise and reproducible placement documentation. EMG-BIDS is now part of BIDS as of version 1.11.0, supported by existing tools, including MNE-BIDS and EEGLAB. We demonstrate the specification through public datasets, including high-density surface EMG recordings. EMG-BIDS provides the foundation for FAIR (Findable, Accessible, Interoperable, Reusable) EMG data sharing, enabling meta-analyses, multi-site studies, and machine learning applications that require standardized, well-documented datasets.

## 1 Introduction

Electromyography (EMG) records the electrical activity produced by skeletal muscles, providing a window into the neuromuscular system that has proven invaluable across clinical medicine, rehabilitation, ergonomics, sports science, neuroscience, and brain-computer interfaces. Since the first human EMG recordings in the early 20th century, the technique has evolved from single-channel needle recordings [1] to sophisticated multi-channel surface arrays that capture muscle activity at high spatiotemporal resolution [2].

While invasive EMG (iEMG) is a fundamental tool in clinical neurology [3], surface EMG (sEMG) offers several practical advantages that have driven its widespread adoption: it is non-invasive, relatively inexpensive, and can be applied due to diverse environments from clinical laboratories to real-world settings. High-density surface EMG (HD-sEMG) systems with electrode arrays of tens to several hundred channels enable spatial mapping of muscle activity, estimation of muscle fiber conduction velocity, and decomposition into individual motor unit action potentials [4]. This technological evolution has expanded EMG applications from basic amplitude measurements to sophisticated analyses, including motor unit tracking, muscle fiber conduction velocity estimation, and fatigue monitoring.

Despite these technical advances, EMG research faces a critical challenge: the absence of standardized data formats and metadata conventions. Unlike neuroimaging modalities such as MRI, where community standards have long facilitated data sharing, EMG data remain fragmented across proprietary formats, undocumented electrode placements, and inconsistent metadata practices. This heterogeneity impedes reusability and reproducibility, prevents large-scale data aggregation, and limits the development of generalizable analysis methods and machine learning models.

The electrode placement problem is particularly acute for EMG. Variations in sensor positioning (2–3 cm) can produce amplitude differences of up to 50% [5], yet most shared datasets provide very limited documentation of electrode placement. Guidelines such as SENIAM (Surface EMG for Non-Invasive Assessment of Muscles) [6] and CEDE (Consensus for Experimental Design in Electromyography) [7] provide recommendations for conducting and reporting EMG measurements, but no standard exists for encoding placement information in a machine-readable and interoperable format that accompanies the data.

### 1.1 The Brain Imaging Data Structure

The Brain Imaging Data Structure (BIDS) emerged as a community-driven effort to standardize data organization and documentation for neuroimaging research [8, 9]. BIDS specifies a cohesive directory structure and file naming convention for the data, alongside JSON and TSV sidecar files that capture essential metadata, guiding researchers not only in how to organize their data but also in what information to document and share. The standard has been adopted by notable data archives (such as OpenNeuro, the NIMH Data Archive, NITRC, publicnEUro, Austrian NeuroCloud, NEMAR, Baby Open Brains) and is supported by an extensive ecosystem of tools.

Although initially conceived for MRI data, BIDS has been extended to multiple modalities: MEG-BIDS for magnetoencephalography [10], EEG-BIDS for electroencephalography [11], iEEG-BIDS for intracranial electrophysiology [12], and Motion-BIDS for motion capture data [13], among others. Each extension preserves the core BIDS principles while addressing modality-specific requirements. Underlying all of these is the FAIR framework (Findability, Accessibility, Interoperability, and Reusability) [14]: by adopting community standards, researchers ensure their data can be discovered, accessed, integrated with other datasets, and reused for purposes beyond the original study.

### 1.2 Scope of EMG-BIDS

EMG-BIDS specifications extend BIDS to support electromyographic recordings from any acquisition system, and is part of the BIDS specifications as of version 1.11.0. The development process entailed an original Github issue^1^ to the BIDS specifications, regular meetings with stakeholders to discuss necessary elements for adequate description of EMG recordings, creating a suite of examples, obtaining a BIDS Extension Proposal Number (BEP044), working on the pull request^2^ and extending the BIDS examples and schema to accommodate the new structures required for the specification. The resulting EMG-BIDS specification covers:

- Surface EMG (sEMG), including bipolar and monopolar configurations
- High-density surface EMG with electrode arrays and grids
- Simultaneous recordings from multiple anatomical locations
- Synchronized recordings from multiple devices

EMG-BIDS focuses on raw data organization and does not prescribe specific analysis pipelines or processing steps. Derived data (e.g., decomposed motor units, force predictions, etc.) fall under the BIDS Derivatives specifications and proposals.

Here, we describe the initial implementation of EMG-BIDS as of BIDS version 1.11.0. The specifications may be adjusted and extended over time to accommodate the emergent community needs and data modalities such as file formats with 32-bit resolution, and inclusion of intramuscular recording. For the most up to date descriptions and best practices consult the BIDS documentation (https://bids-specification.readthedocs.io).

## 2 EMG-BIDS Specification

### 2.1 General Principles and File Structure

EMG-BIDS follows the core BIDS directory hierarchy. Data are organized by subject and optionally by session, with EMG files stored in a dedicated `emg/` subdirectory (Figure 1). File names follow the BIDS entity-based convention:

**Fig. 1.**
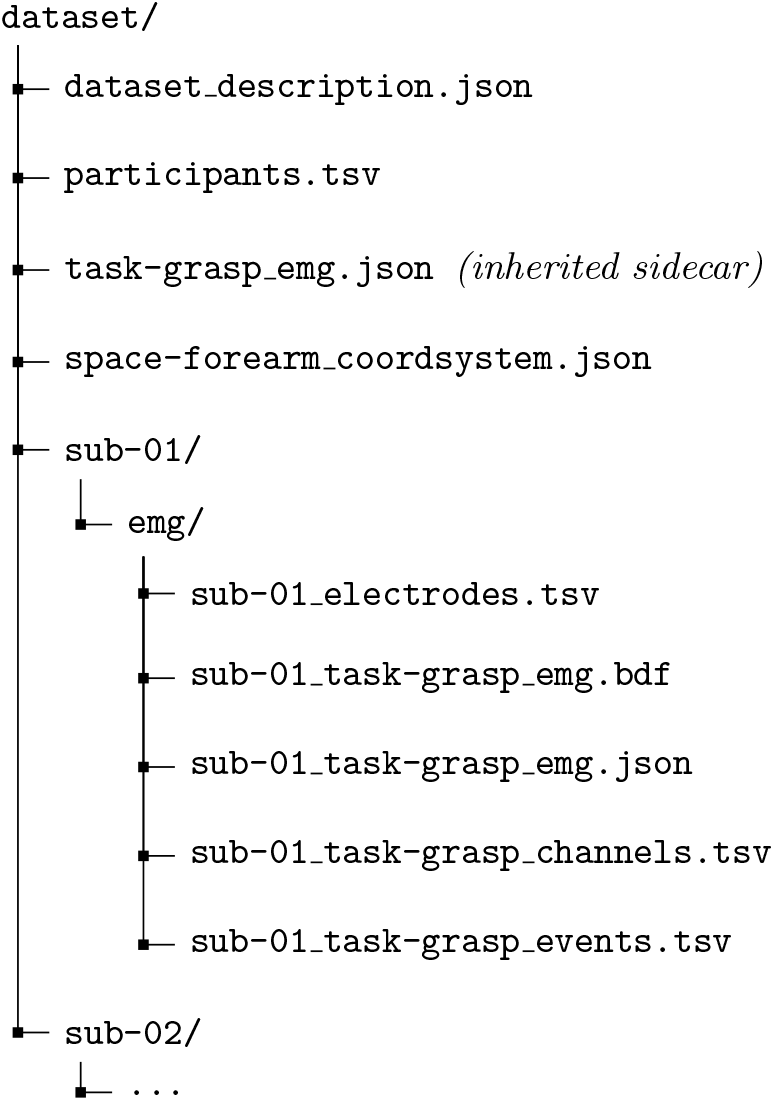
EMG-BIDS directory structure. Data are organized by subject with EMG files in a dedicated `emg/` subdirectory. Each recording includes the data file (`.bdf` or `.edf`), a JSON sidecar with acquisition metadata, channel descriptions (`channels.tsv`), electrode positions (`electrodes.tsv`), and event markers (`events.tsv`). Root-level JSON files provide inherited metadata for all recordings.

~~~
sub-<label>[_ses-<label>]_task-<label>
    [_acq-<label>][_run-<index>]
    [_recording-<label>]_emg.<extension>
~~~

The `task` entity is required and indicates the experimental condition or activity during which data were recorded. The `recording` entity distinguishes data from multiple EMG devices recording simultaneously into separate files.

### 2.2 Data Formats

EMG-BIDS supports two data formats: the European Data Format (EDF/EDF+) and the BioSemi Data Format (BDF/BDF+). Both formats are open, well-documented, and supported by major analysis packages. BDF+ is recommended over EDF+ due to its higher resolution (24-bits vs. 16-bits per sample) and better support for event metadata. Both formats can store multiple channels with heterogeneous sampling frequencies, accommodating systems that record EMG alongside auxiliary signals (accelerometers, force sensors) at different rates.

### 2.3 Metadata Sidecars

Each EMG data file is accompanied by a JSON sidecar (`*_emg.json`) containing acquisition parameters and experimental details. Table 1 summarizes the EMG-specific metadata fields.

**Table 1.** EMG-BIDS JSON sidecar fields. Requirement levels follow RFC 2119 conventions as adopted by BIDS: REQUIRED fields must be present; RECOMMENDED fields should be included when available.

| Field | Level | Description |
| --- | --- | --- |
| SamplingFrequency | REQUIRED | Sampling rate in Hz |
| PowerLineFrequency | REQUIRED | Power line frequency (50 or 60 Hz) |
| EMGPlacementScheme | REQUIRED | How electrode locations were determined: "Measured", "ChannelSpecific", or "Other" |
| EMGReference | REQUIRED | Reference electrode specification or "Bipolar" for paired sensors |
| RecordingType | REQUIRED | Data continuity: "continuous", "epoched", or "discontinuous" |
| SoftwareFilters | REQUIRED | Digital filters applied, or "n/a" |
| EMGPlacementSchemeDescription | REQUIRED* | Description when EMGPlacementScheme is "Other" |
| EMGChannelCount | RECOMMENDED | Number of EMG channels |
| HardwareFilters | RECOMMENDED | Analog filter specifications |
| TaskName | RECOMMENDED | Short task identifier |
| TaskDescription | RECOMMENDED | Detailed task description |
| Manufacturer | RECOMMENDED | Recording system manufacturer |
| EMGGround | OPTIONAL | Ground electrode location |
| InterelectrodeDistance | OPTIONAL | Distance between electrode pairs (mm) |
| SkinPreparation | OPTIONAL | Skin preparation procedure |

\*REQUIRED only if EMGPlacementScheme is marked as "Other"

The `EMGPlacementScheme` field addresses the critical need for placement transparency. When set to `“Measured”`, electrode coordinates are provided in accompanying tabular files. When `“Channel Specific”`, placement details vary across sensors and are documented per-channel. When `“Other”`, a free-text description in `EMGPlacementSchemeDescription` explains the procedure (e.g., “SENIAM guidelines for tibialis anterior”).

BIDS inheritance allows common metadata to be specified once at higher directory levels, reducing redundancy. A root-level `emg.json` file can define equipment parameters shared across all recordings, with acquisition-specific values (e.g., `RecordingDuration`) in per-file sidecars.

### 2.4 Channel Description

The `*_channels.tsv` file provides per-channel metadata (Table 2). Unlike EEG where channels typically correspond one-to-one with electrodes, EMG channels often represent differential recordings between electrode pairs or signals from multi-electrode sensors.

**Table 2.** Columns in *_channels.tsv. The signal_electrode column links channels to physical electrodes, enabling disambiguation of complex recording configurations.

| Column | Level | Description |
| --- | --- | --- |
| name | REQUIRED | Channel identifier matching data file |
| type | REQUIRED | Channel type (EMG, ECG, REF, etc.) |
| units | REQUIRED | Physical units (e.g., $\mu\text{V}$, mV) |
| signal_electrode | RECOMMENDED | Name of signal electrode in `*_electrodes.tsv` |
| reference | RECOMMENDED | Reference electrode name |
| target_muscle | RECOMMENDED | Anatomical muscle name |
| group | RECOMMENDED | Device or array identifier |
| sampling_frequency | OPTIONAL | If different from rate specified in sidecar file |

The `units` column addresses a common challenge in EMG: different systems report signals in different units (V, mV, *µ*V) with different gain stages. Explicit unit specification ensures correct interpretation regardless of acquisition system.

### 2.5 Electrode Description and Coordinate Systems

The `*_electrodes.tsv` file lists physical electrodes with their spatial coordinates (Table 3). The `x`, `y`, and (optionally) `z` columns contain numeric position values whose meaning is defined by the associated coordinate system file (`space-<label> coordsystem.json`). For example, coordinates may represent millimeters relative to the corner of an electrode grid, or percentages along an anatomical axis between two landmarks. Unlike the original BIDS electrode specification designed for scalp EEG, EMG-BIDS accommodates electrodes placed at multiple anatomical locations; a forearm array, leg sensors, and facial electrodes might all be recorded simultaneously.

**Table 3.**
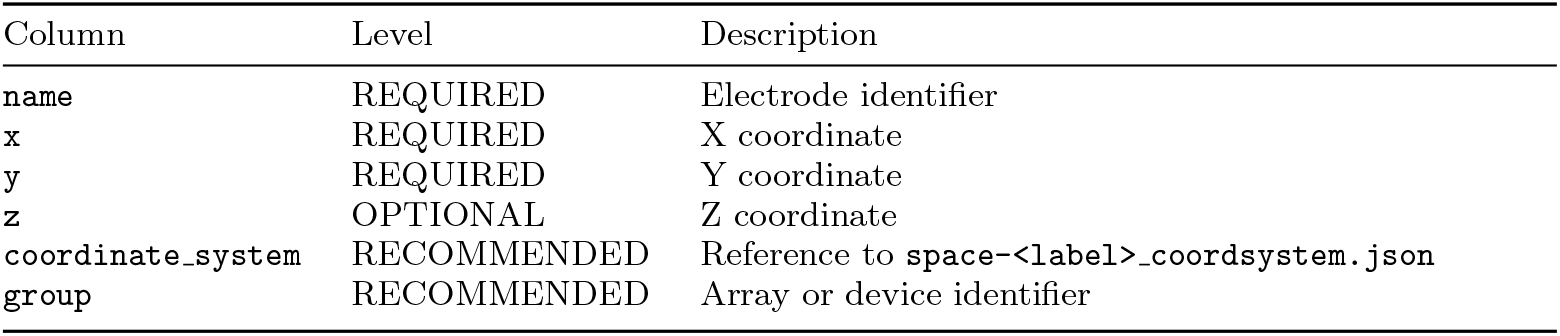
Columns in *_electrodes.tsv. The coordinate system column enables electrodes from different body regions to coexist, each referenced to an appropriate anatomical coordinate frame.

| Column | Level | Description |
| --- | --- | --- |
| `name` | REQUIRED | Electrode identifier |
| `x` | REQUIRED | X coordinate |
| `y` | REQUIRED | Y coordinate |
| `z` | OPTIONAL | Z coordinate |
| `coordinate_system` | RECOMMENDED | Reference to `space-<label>_coordsystem.json` |
| `group` | RECOMMENDED | Array or device identifier |

Each coordinate system is defined in a corresponding `space-<label>_coordsystem.json` file specifying the anatomical axes and units. EMG-BIDS introduces *hierarchical coordinate systems* to handle high-density electrode arrays: a child coordinate system (such as an electrode grid in millimeters) is anchored to a parent anatomical coordinate system (such as using normalized coordinates), making placement transferable across subjects with different body proportions. Figure 3 illustrates this hierarchy for the multi-device recording in Figure 2, where HD-sEMG grids on the vastus lateralis (VL) and vastus medialis (VM) are each defined in local millimeter coordinates and anchored to a parent thigh system defined by palpable bony landmarks.

**Fig. 2.**
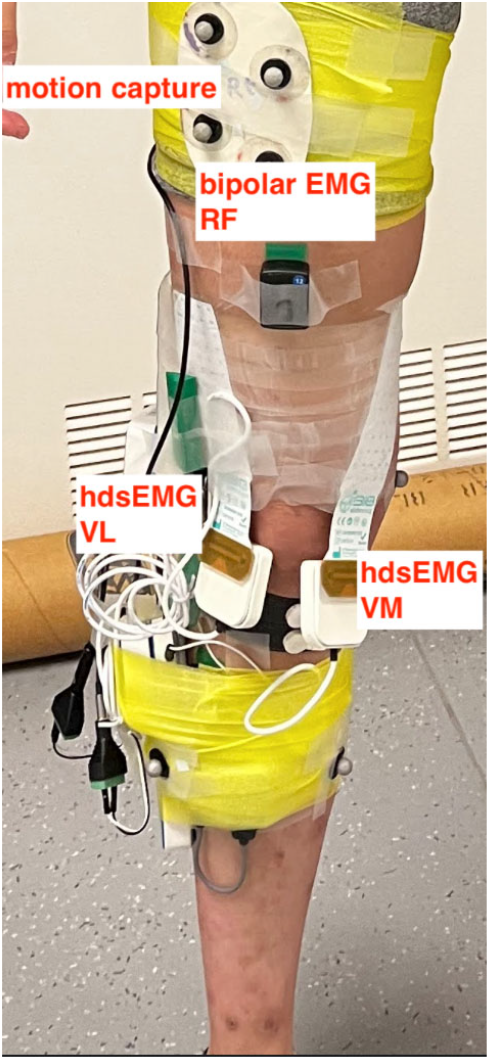
Multi-device EMG recording from the `emg_ConcurrentIndependentUnits` example (see github.com/bids-standard/bids-examples). The setup includes motion capture markers, a bipolar EMG sensor on the rectus femoris (RF), and two HD-sEMG arrays on the vastus lateralis (VL) and vastus medialis (VM). EMG-BIDS accommodates such configurations through multiple coordinate systems, the `group` column linking channels to devices, and hierarchical coordinate systems anchoring local grids to anatomical landmarks.

**Fig. 3.**
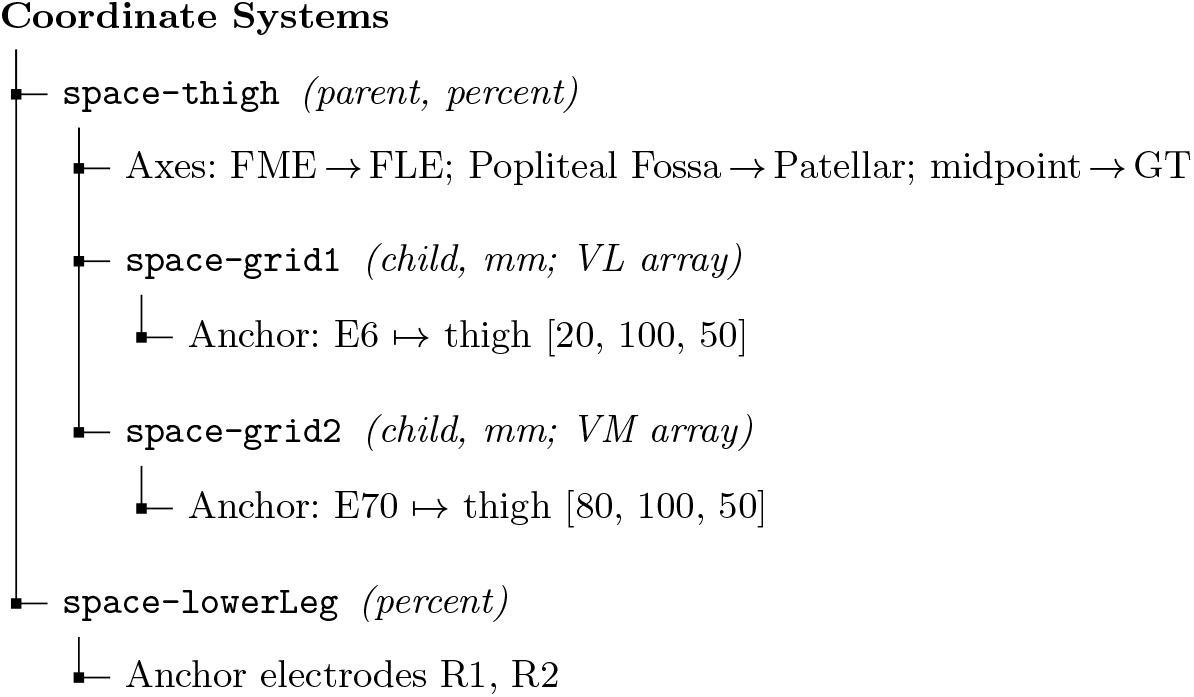
Hierarchical coordinate system structure for the recording in Figure 2. The parent `thigh` system uses anatomical landmarks (Femur Medial Epicondyle (FME), Femur Lateral Epicondyle (FLE), Greater Trochanter (GT)) with normalized coordinates in percent. HD-sEMG grids, placed on Vastus Medialis (VM) and Vastus Lateralis (VL) muscles, are defined in a child system (in mm) anchored to the parent via `AnchorElectrode` and `AnchorCoordinates`. A separate `lowerLeg` system locates the anchor electrodes.

This hierarchical approach, inspired by the Unified Sensor Placement Framework [15], enables precise documentation of electrode positions within a local sensor grid while situating the position of that grid with respect to reproducible anatomical landmarks. Figure 2 depicts the exemplar structure of the multi-space coordinate systems with proper anchoring of child coordinate systems (HD-sEMG grids) to the parent (thigh); Table 4 shows how this information is conveyed in an `*_electrodes.tsv` file, by providing coordinate system names alongside the coordinate values on each row. The link between child and parent coordinate systems is made explicit within each child `*_coordsystem.json` file using the keys `ParentCoordinateSystem`, `AnchorElectrode`, and `AnchorCoordinates`.

**Table 4.** Excerpt from *_electrodes.tsv for the HD-sEMG recording in Figure 2. Grid electrodes reference their child coordinate systems (in mm), while anchor electrodes (reference electrodes used to link a child coordinate system to its parent) are in the `lowerLeg` system (in percent). The bipolar RF sensor uses the common SENIAM-based workflow: placement documented via `EMGPlacementScheme` and `target muscle` in `*_channels.tsv`.

| name | x | y | coordinate_system | group |
| --- | --- | --- | --- | --- |
| E1 | 0 | 0 | grid1 | Grid1 |
| E2 | 0 | 8 | grid1 | Grid1 |
| ... (64 electrodes, 8 mm inter-electrode distance) |  |  |  |  |
| E65 | 0 | 0 | grid2 | Grid2 |
| E66 | 0 | 8 | grid2 | Grid2 |
| ... (64 electrodes, 8 mm inter-electrode distance) |  |  |  |  |
| R1 | 100 | 50 | lowerLeg | Grid1 |
| R2 | 0 | 50 | lowerLeg | Grid2 |

For bipolar sensors where two electrodes are rigidly attached as a unit and a single (differential) signal is recorded from the device, the `EMGReference` field is set to `“Bipolar”`. EMG-BIDS supports several approaches for documenting the placement of such sensors, reflecting the range of practices in the field:

1. **Guideline-based placement (most common):** Researchers following established protocols such as SENIAM set `EMGPlacementScheme` to `“Other”` and describe the protocol in `EMGPlacementSchemeDescription` (e.g., “SENIAM guidelines for tibialis anterior”). The target muscle is documented per-channel in `*_channels.tsv`. No digitized coordinates are required. The `emg_Independent Mod` example in the BIDS examples repository illustrates this workflow.
2. **Channel-specific descriptions:** When placement varies across sensors and does not follow a single protocol, `EMGPlacementScheme` is set to `“ChannelSpecific”` and per-channel columns in `*_channels.tsv` (such as `placement_scheme` and `placement_description`) document each sensor location individually.
3. **Measured coordinates:** When electrode positions are digitized, `EMGPlacementScheme` is set to `“Measured”` and coordinates for each pole (or the sensor centroid, depending on available information) are provided in `electrodes.tsv` with an associated coordinate system file.

The bipolar RF sensor in Figure 2 follows the first approach, using a guideline-based description without digitized coordinates, while the HD-sEMG grids in the same recording use measured coordinates with hierarchical coordinate systems.

### 2.6 Multi-Device and Multi-Unit Support

EMG recordings increasingly involve multiple recording units, either connected to a synchronized recording hub (such as multiple bipolar EMG recording modules wirelessly connected to a hub), or as independent modules synchronized via TTL triggers or synchronization platforms [16], such as a high-density sEMG array on one muscle, bipolar sensors on another, and anchor electrodes elsewhere. EMG-BIDS supports this through:

- The `recording` entity in file names to separate data acquired simultaneously from different devices
- The `group` column in `*_channels.tsv` and `*_electrodes.tsv` to identify channels and electrodes within a recording device.
- Per-channel `units` to accommodate systems with different gain/scale configurations
- Multiple coordinate system files for electrodes at different anatomical locations

## 3 Example Datasets

Eight schematic EMG datasets (with empty data files but otherwise exhibiting valid BIDS format) are available in the official BIDS examples repository (https://github.com/bids-standard/bids-examples). Table 5 summarizes these examples, each illustrating specific features of the specification. In addition, two full-scale converted datasets are described below.

**Table 5.** EMG-BIDS example datasets in the BIDS examples repository. Each example demonstrates a different recording configuration and set of specification features. Note that these datasets are schematic with empty data files.

| Example | Description |
| --- | --- |
| `emg_CustomBipolar` | Single custom bipolar sensor on forearm flexors |
| `emg_CustomBipolarFace` | Several bipolar sensors on facial muscles during speech |
| `emg_IndependentMod` | Wireless bipolar modules on multiple arm muscles |
| `emg_TwoWristbands` | Two dry-electrode wristbands (bilateral forearm) with coordinate systems |
| `emg_TwoHDsEMG` | Two HD-sEMG grids with hierarchical coordinate systems |
| `emg_MultiBodyParts` | Multiple electrode types (surface, fine-wire, needle) across body regions |
| `emg_ConcurrentIndependentUnits` | Concurrent bipolar and HD-sEMG at different sampling rates (Figure 2) |
| `emg_Multimodal` | EMG integrated with EEG and motion capture |

### 3.1 Hyser High-Density Surface EMG Dataset

The Hyser (High densitY Surface Electromyogram Recordings) dataset [17] includes 20 subjects performing 5 task types (pattern recognition gestures, maximum voluntary contraction, single-finger and multi-finger force tracking, and random combinations) recorded with 256-channel HD-sEMG from four 8× 8 electrode arrays on the forearm. Hyser dataset was originally shared in a custom structure on PhysioNet [18]. We transformed this dataset to EMG-BIDS (Hyser-BIDS). Hyser-BIDS dataset includes:

- Hierarchical coordinate systems: four child grids (ExtensorDistal, ExtensorProximal, FlexorDistal, FlexorProximal) anchored to a parent forearm system
- Complete channel-to-electrode mapping via `signal_electrode`
- Event files with gesture timing and trial structure
- Synchronized force data as BIDS physio files

We verified data integrity through correlation analysis between original WFDB files and converted BDF, achieving mean correlations exceeding 0.9999 across all recordings. Hyser-BIDS dataset is available on NEMAR as `nm000108` [19].

### 3.2 EMG2Qwerty Typing Dataset

The emg2qwerty dataset [20] is a large-scale forearm EMG dataset recorded during keyboard typing, comprising 1,135 sessions from 108 participants and 346 hours of recording. Each participant wore two 16-channel dry-electrode wristbands while typing prompted text on a standard keyboard. The dataset includes bilateral EMG signals, keystroke events with exact timing and key labels, and session metadata.

The EMG-BIDS conversion preserves the complete keystroke event structure, bilateral wristband recordings using the `recording` entity to separate left and right devices, and registers approximate bipolar channel locations. The converted dataset is available on NEMAR as `nm000104` [21].

### 3.3 MUniverse Decomposition Benchmark Collection

The MUniverse motor unit decomposition benchmarking suite [22] includes 3 experimental and 3 simulated datasets containing a total of 11,230 HD-sEMG recordings (50 to 320 electrodes) from different anatomical regions (lower leg, upper leg, and forearm) and considering diverse isometric, ballistic, and dynamic tasks. The original experimental datasets have been converted from custom formats to EMG-BIDS and are available on Harvard Dataverse. The focus of this data collection is on large-scale fully-automated computer-assisted data analysis, i.e., the decoding of motor unit activity from HD-sEMG data.

## 4 Community Tools and Infrastructure

### 4.1 Software Support

EMG-BIDS is supported by existing BIDS tools with minimal or no modification required:

- **MNE-BIDS [23, 24]**: Python package for reading and writing BIDS electrophysiology data, including EMG
- **EEGLAB [25]**: MATLAB toolbox with `EEG-BIDS` plugin for BIDS export
- **FieldTrip [26]**: MATLAB toolbox supporting BIDS data organization
- **bids-validator [27]**: Command-line and web-based validation of BIDS datasets as of version 2.4.0
- **MUniverse [22]**: Python package for benchmarking motor unit identification algorithms, that includes a module for reading and writing EMG-BIDS data

The EDF/BDF formats used by EMG-BIDS are readable by standard libraries including pyedflib and edfio (Python), edfread (MATLAB), and EDFlib (C).

## 5 Discussion

EMG-BIDS addresses long-standing challenges in EMG data sharing by providing a community standard compatible with existing BIDS infrastructure. Several design decisions merit discussion.

### 5.1 Electrode Placement Transparency

The most significant innovation in EMG-BIDS is the framework for documenting electrode placement. Unlike EEG, where coordinate system axes are almost universally defined with respect to the same set of anatomical landmarks (pre-auricular points, nasion, and inion), EMG electrodes can be placed anywhere on the body, and variation across subjects can make some approaches to electrode placement hard to replicate. The `EMGPlacementScheme` field accommodates the most common approaches to electrode placement (visual, palpated, or measured proximity to landmarks; functional localization), while support for digitized electrode coordinates and hierarchical coordinate systems provides the flexibility to document modern high-density array devices easily as well.

For high-density arrays, the local grid coordinates capture the physical layout of the sensor array, while the anchor position and parent coordinate system enable replicability across subjects and studies. For individual electrodes, the parent system alone may suffice; the complexity of nested coordinate systems is available if needed but not imposed on datasets that don’t require it. This design also aligns with the Unified Sensor Placement Framework [15], which proposes standardized anatomical coordinate systems across modalities.

### 5.2 Relationship to Existing Standards

There are valuable standards for conducting and reporting EMG experiments; however, these guidelines are typically developed for human-readable documentation rather than machine-readable metadata, for example, the SENIAM project [6] or those released by the International Society of Electromyography and Kinesiology [7, 28, 29]. EMG-BIDS builds on top of these guidelines by providing well-defined metadata fields for all required items. Researchers can also include additional custom metadata fields in their JSON sidecars and TSV files as needed; BIDS is designed to be extensible without breaking compatibility with existing tools. SENIAM guidelines remain particularly valuable for specific pre-defined muscle placements. In detail, researchers following SENIAM guidelines should set `EMGPlacementScheme` to `“Other”` and describe their procedure in `EMGPlacementSchemeDescription`, or provide measured coordinates if digitization was performed.

### 5.3 Limitations and Future Directions

#### 5.3.1 Coordinate Systems

EMG-BIDS currently supports only `“Other”` for the `EMGCoordinateSystem` field, as standardized coordinate system names for body segments (other than the head/skull) have not yet been established. Future versions may incorporate the anatomical coordinate systems from the Unified Sensor Placement Framework or some other standard, once community consensus is reached.

#### 5.3.2 Derivatives

The specification focuses on raw data. Standards for EMG derivatives (decomposed motor units, normalized envelopes, fatigue indices) will be addressed through BIDS-Derivatives as these analysis methods mature.

#### 5.3.3 HED Integration

Integration with the Hierarchical Event Descriptors (HED) system would enable richer annotation of EMG events, particularly for complex experimental protocols. This represents an area for future development.

## 6 Conclusion

EMG-BIDS extends the Brain Imaging Data Structure to electromyography, providing a community standard for organizing EMG data that addresses the unique challenges of this modality: diverse tasks, heterogeneous electrode types and configurations, critical dependence on electrode placement, and diverse anatomical regions. The hierarchical coordinate system framework enables transparent documentation of sensor positions from individual electrodes to high-density arrays, laying the foundation for reproducible EMG research.

EMG-BIDS is now part of BIDS version 1.11.0, supported by existing tools and data repositories. By adopting EMG-BIDS, researchers contribute to a growing ecosystem of standardized, FAIR neurophysiology data that enables meta-analyses, multisite collaborations, and machine learning applications requiring large, well-documented datasets.

We invite the EMG research community to adopt EMG-BIDS for data sharing and to contribute to its continued development through the open BIDS specification process.

## Acknowledgements

We thank the BIDS community for feedback during the extension proposal process, and the developers of MNE-BIDS and EEGLAB for tool support.

## Declarations

- **Funding**: This work was partially supported by the NIH R01NS047293 to AD. SYS, DM, EL, and AD thank Meta Reality Labs for unrestricted gifts that helped with timely turnaround of this specification.
- **Competing interests**: The authors declare no competing interests.
- **Data availability**: Converted datasets are available on NEMAR (https://nemar.org): Hyser (nm000108) and emg2qwerty (nm000104). The original Hyser data are at PhysioNet (doi:10.13026/ym7v-bh53). BIDS example datasets are at https://github.com/bids-standard/bids-examples.
- **Code availability**: The BIDS specification is available at https://bids-specification.readthedocs.io. Conversion tools are available through MNE-BIDS and EEGLAB.
- **Author contributions**: SYS conceived and proposed the EMG-BIDS standard. All authors contributed to the specification development. DM drafted the initial pull request and implemented the BIDS schema. RG and CJM coordinated alignment with the BIDS standard as BIDS maintainers. SYS wrote the manuscript with contributions from all authors. All authors reviewed and approved the final manuscript.

## Footnotes

1 https://github.com/bids-standard/bids-specification/issues/1371

2 https://github.com/bids-standard/bids-specification/pull/1998

